# Anle138b manifests potent anti-hyperglycemic activity in type 2 diabetic mice expressing hIAPP

**DOI:** 10.1101/2024.08.27.609850

**Authors:** Mohammed M. H. Albariqi, Sanne M.G. Baauw, Sjors J.P.J. Fens, Sabine Versteeg, Sergey Ryazanov, Andrei Leonov, Hanneke L.D.M. Willemen, Nikolas Stathonikos, Raina Marie Seychell, Adam El Saghir, Bram Gerritsen, Lucie Khemtemourian, Neville Vassallo, Armin Giese, Niels Eijkelkamp, Christian Griesinger, Jo W. M. Höppener

## Abstract

**Aims/Hypothesis:** Type 2 diabetes mellitus (T2DM) is a common metabolic disease, characterized by impaired insulin action, often called “insulin resistance”, in conjunction with a decline in insulin secretion by the pancreatic islet beta cells, referred to as “beta cell failure”. Both conditions contribute to causing hyperglycemia, which is the central feature of T2DM. Cytotoxic aggregates of the beta cell hormone human islet amyloid polypeptide (hIAPP), promote T2DM pathogenesis by damaging the pancreatic islet beta cells and reducing insulin secretion.

Anle138b is an amyloid oligomer modulator with demonstrated disease-modifying properties in mouse models of neurodegenerative diseases linked to protein aggregation, and is currently in a phase 2 clinical trial. We therefore tested whether anle138b can reduce hyperglycemia in a severe hIAPP transgenic mouse model of T2DM.

**Methods:** Anle138b was administered orally via dietary admixture to male *hIAPP Ob/Ob* mice and *Ob/Ob* controls; blood glucose-, insulin-, and glycated hemoglobin (HbA1c) levels were measured at different timepoints over 34 weeks; in addition, the effects of anle138b on the histomorphology of the pancreatic islets was determined. Finally, anle138b effects on mitochondrial toxicity of hIAPP aggregates was assessed *in vitro*.

**Results:** Oral administration of anle138b in male *hIAPP Ob/Ob* mice potently reduced hyperglycemia and glycated HbA1c levels compared to non-treated mice (at 34 weeks, mean glucose level non-treated: 30 ± 1.3 mM, treated: 15 ± 2.2 mM (*p* < 0.001); at 32 weeks, mean HbA1c level non-treated: 77 ± 3.3 mM/M, treated: 41 ± 5.3 mM/M (*p* < 0.01)), in a hIAPP-dependent way. In addition, anle138b increased both the number of pancreatic islets as well as the overall islet beta cell mass (by 54% and 58%, respectively) in *hIAPP Ob/Ob* mice. *In vitro*, anle138b inhibited toxic effects of hIAPP on mitochondria, such as mitochondrial swelling (decreased by 92%, *p* < 0.01), release of cytochrome *c* (decreased by 63.7%, *p* < 0.001) and lowered mitochondrial membrane potential (increased by 62%, *p* < 0.001).

**Conclusions/Interpretation:** Collectively, our results indicate that anle138b is a promising clinical drug candidate for preserving islet function and reducing hyperglycemia in T2DM, in part by mitigating mitochondrial dysfunction.

**Research in Context:** *What is already known about this subject?:*

- Type 2 diabetes mellitus (T2DM) is characterized by hyperglycemia, which stems from impaired insulin action coupled with declining insulin secretion from pancreatic islet beta cells.
- Aggregation of the beta cell hormone human islet amyloid polypeptide (hIAPP) plays a crucial role in the loss and dysfunction of the insulin-secreting beta cells.
- The small-molecule anle138b effectively modulates the aggregation of amyloidogenic proteins in neurodegenerative conditions such as Alzheimer’s and Parkinson’s disease.

*What is the key question?:*

- Can the oligomer modulator anle138b ameliorate the diabetic phenotype in a severe hIAPP transgenic mouse model of T2DM (*hIAPP Ob/Ob)* ?
- What are the new findings?
- Oral treatment of *hIAPP Ob/Ob* mice with anle138b causes reduced hyperglycemia and reduced glycated hemoglobin (HbA1c) levels, in a hIAPP-dependent manner.
- Anle138b increases pancreatic islet number and beta cell mass and restores the inverse relation between insulin and glucose in *hIAPP Ob/Ob* mice.
- *In vitro* anle138b inhibits toxic effects of hIAPP aggregates on mitochondria.

**How might this impact on clinical practice in the foreseeable future?:**

- Anle138b could be introduced as a promising clinical drug candidate for slowing down disease progression of T2DM.

## 1. Introduction

Type 2 diabetes mellitus (T2DM) is a common metabolic disease, affecting more than 500 million people globally in 2021 and this number is expected to rise to more than 700 million by 2045 [1]. The WHO has declared T2DM as the first non-infectious epidemic [2]. Because T2DM is strongly associated with obesity, these two conditions are together also referred to as the “twin epidemics” [3]. T2DM is characterized by impaired insulin action, often called “insulin resistance”, and by insufficient insulin production by the pancreatic islet β cells, referred to as “beta cell failure”, causing hyperglycemia which is the central feature of T2DM [4].

A distinct histopathological feature of T2DM is the presence of extracellular amyloid (fibrillar, congophilic protein deposits) within the pancreatic islets. This “islet amyloid” is detected in 80-90% of T2DM patients and is mainly composed of the hormone islet amyloid polypeptide (IAPP), also named amylin [5,6]. IAPP is co-produced and co-secreted with insulin from the pancreatic islet beta cells. As a monomer, IAPP is a soluble protein involved as a hormone in, amongst others, gastric emptying and satiety; but when overproduced, particularly under conditions of insulin resistance, IAPP can aggregate and deposit as amyloid. In humans, monkeys and cats with T2DM, islet amyloid formation has been associated with beta cell failure [7]. Conversely, in mice and rats, species which do not naturally develop T2DM, IAPP does not form amyloid due to a different amino acid sequence [8]. Amyloid fibrils feature a characteristic cross-β structure, formed by stacking of β-sheets from the fibril-forming protein [9]. Soluble pre-fibrillar human IAPP (hIAPP) oligomers, that form in the lag phase of amyloid fibril formation, have been shown to be the most harmful species of hIAPP, and are directly toxic to insulin-producing islet beta cells [10]. Increasing evidence shows that hIAPP can cause mitochondrial dysfunction [11] and soluble toxic hIAPP oligomers significantly contribute to beta cell dysfunction and metabolic dysregulation associated with T2DM [12]. Transgenic rodent models that overexpress hIAPP in islet beta cells have been shown to develop islet amyloid and display impaired glucose homeostasis [13–15]. In such hIAPP transgenic mice, islet amyloid develops in both sexes, but generally more frequently in males than in females [13–14].

More broadly, protein aggregation and its associated cytotoxicity are linked to various prevalent and debilitating chronic diseases, such as Alzheimer’s disease (AD), Parkinson’s disease (PD) and prion diseases. A growing body of literature indicates that hIAPP aggregation in pancreatic islets plays a similar role in T2DM as the amyloid-beta, tau and α-synuclein proteins do for the neurological aggregation diseases AD and PD [16]. Therefore, small molecule compounds and antibodies that interfere with amyloid protein aggregation are being developed to reduce amyloid-associated cell and tissue damage in these diseases. This is exemplified by the recent success of lecanemab, a humanized monoclonal antibody against diffusible aggregates of the AD-associated amyloid-beta protein [17], and antibodies against hIAPP proto-fibrils [18]. In addition, active immunization against hIAPP oligomers showed ameliorating effects in a transgenic T2DM mouse model [19].

Despite the pathogenic role of hIAPP in T2DM, therapies targeting hIAPP aggregation are currently an unmet need for T2DM patients, probably in part because the role of hIAPP aggregation as a main driver of T2DM pathogenesis is still in the early stages of clinical research [16]. The diphenyl-pyrazole (DPP) compound anle138b modulates early aggregation of alpha-synuclein, the key protein in the pathogenesis of PD, and inhibits aggregation of α-synuclein *in vivo* after oral administration in transgenic PD mouse models [20,21]. Importantly, anle138b reduced disease progression in mouse models for the neurodegenerative amyloid disorders AD (based on aggregation of tau) [22,23], PD [20] multiple system atrophy (MSA) [24] and prion diseases [20].

Anle138b has been tested in a phase Ia study (NCT04208152) and in a phase Ib clinical trial (NCT04685265), where it showed no adverse effects at concentrations expected to be therapeutically efficacious [25]; it is currently in a phase 2 clinical study on MSA (NCT06568237). Since anle138b was effective in mouse models of various neurodegenerative proteinopathies, by interfering with aggregation of α-synuclein, amyloid-beta, tau and prion protein, we hypothesized that anle138b may also have disease-modifying effects for T2DM by acting on hIAPP, hence lowering blood glucose and blood HbA1c levels. To test this notion, the therapeutic potential of anle138b was assessed by oral administration of the drug via dietary admixture to transgenic obese mice overexpressing hIAPP in their pancreatic islet beta cells (*hIAPP Ob/Ob*) [14]. Contrary to non-obese *hIAPP* mice [26], the *hIAPP Ob/Ob* mice develop severe insulin resistance, hypoinsulinemia and marked hyperglycemia already at a young age, and this is associated with deposition of amyloid in their pancreatic islets [14]. In addition to measurement of glucose-, HbA1c- and insulin levels, the amyloid load of the pancreatic islets as well as islet number and beta cell mass were determined. Finally, effects of anle138b on hIAPP-induced mitochondrial toxicity were assessed *in vitro*.

## 2. Results

### 2.1 Anle138b potently reduces hyperglycemia and glycated HbA1c levels in hIAPP Ob/Ob mice

First we investigated whether anle138b reduces diabetic symptoms in a *hIAPP Ob/Ob* mouse model of T2DM. Therefore, we treated male mice with a dietary admixture containing 2 g anle138b /kg food for 8 months, starting at an age of approximately 3 wk (immediately after weaning), in order to perform preventive or early curative treatment. This dose had also been used in previous studies with other disease models [25–30]. To determine whether possible effects of anle138b treatment are hIAPP-dependent, we also treated non-transgenic *Ob/Ob* mice, which lack hIAPP expression and islet amyloid and which are less severely diabetic [14], with anle138b under the same conditions. Therefore, for both the *hIAPP Ob/Ob* and *Ob/Ob* mice, control groups received the same food but without anle138b.

Remarkably, *hIAPP Ob/Ob* mice fed anle138b had significantly lower blood glucose levels compared to mice fed the same chow diet but without anle138b, after 16 wk and onwards (Fig. 1a,b). At 34 wk, the mean glucose levels of untreated *hIAPP Ob/Ob* mice were 30 ± 1.3 mM, while in the anle138b-treated mice mean glucose levels were 50% less, at 15 ± 2.2 mM (*p* < 0.001). In contrast, anle138b did not affect blood glucose levels in non-transgenic *Ob/Ob* mice, being around 10 mM at 34 wk (Fig. 1a,b). In the *hIAPP Ob/Ob* mice, the blood glucose levels were much lower after overnight fasting as compared to 6 hr of fasting (Fig. 1a *vs.* Fig. 1b). A trend for increased plasma insulin levels was observed after 16 and 23 wk of treatment of *hIAPP Ob/Ob* mice with anle138b, but these differences were not statistically significant (Fig. 1c,d; *p* = 0.35 and 0.07, respectively). Moreover, in *hIAPP Ob/Ob* mice, anle138b treatment did not affect fasting plasma IAPP levels, body weight or food intake (Fig. S1a-c). In *Ob/Ob* mice, anle138b did not affect plasma insulin levels (Fig. 1c,d), plasma IAPP levels, body weight or food intake (Fig. S1a-c).

**Figure 1.**
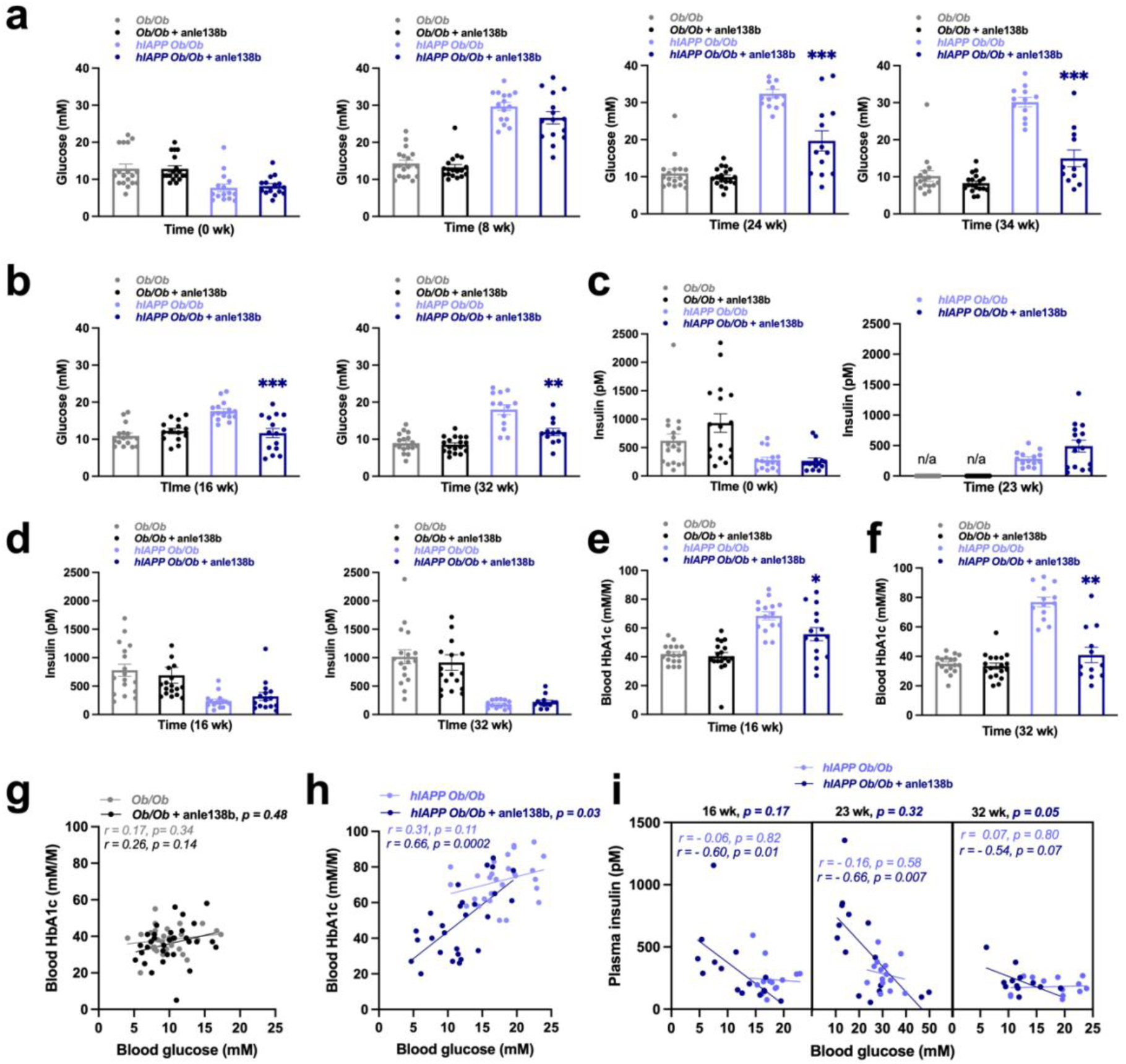
Anle138b reduces hyperglycaemia in *hIAPP Ob/Ob* mice, but not in *Ob/Ob* mice. (**a**) Basal blood glucose levels determined after 4 h fasting in the morning (t=0 wk), or after 6 h fasting in the morning (at 8, 24 and 34 wk) (*n* = 16-18 per group for the *Ob/Ob* mice and *n* = 12-15 per group for the *hIAPP Ob/Ob* mice at each time point). (**b**) Basal blood glucose levels determined after overnight fasting (at 16 and 32 wk) (*n*= 14-18 per group for the *Ob/Ob* mice and *n* = 12-15 per group for the *hIAPP Ob/Ob* mice at each time point). **(c)** Basal plasma insulin levels determined after 4 h fasting in the morning (t=0 wk), or after 6 h fasting in the morning (at 23 wk) (*n* = 18 per group for the *Ob/Ob* mice and *n* = 14-15 per group for the *hIAPP Ob/Ob* mice at each time point). **(d)** Basal plasma insulin levels determined after overnight fasting (at 16 and 32 wk) (*n* = 17-18 per group for the *Ob/Ob* mice and *n* = 13-15 per group for the *hIAPP Ob/Ob* mice at each time po). (**e,f**) Blood HbA1c levels at 16 wk and 32 wk, respectively (*n* = 16-18 per group for the *Ob/Ob* mice and *n* = 12-14 per group for the *hIAPP Ob/Ob* mice at each time point). (**g,h**) Correlation between blood glucose and blood HbA1c levels of *Ob/Ob* and *hIAPP Ob/Ob* mice, respectively (data at 16 and 32 wk combined, *n* = 32 per group for the *Ob/Ob* mice and *n* = 27-28 per group for the *hIAPP Ob/Ob* mice). (**i**) Correlation between blood glucose levels and plasma insulin levels of *hIAPP Ob/Ob* mice treated with anle138b or control diet during 16, 23 and 32 wk (*n*= 12-15 per group at each time point). All experiments were performed in male mice. (**a-f**) Data are presented as individual values, bars indicate mean ± SEM; unpaired two-way Welch’s t-test, \**p* < 0.05, \*\**p* < 0.01, \*\*\**p* < 0.001 for anle138b-treated vs untreated mice. (**g-i)** Pearson Correlation’s test; the correlation coefficient (r) and the two-tailed *p*-value are indicated for both the treated and the untreated mice. Linear regression was performed to examine if the slopes of the two regression lines are significantly different (*p*-value indicated at the top of the figures). Also see Figure S1.

Glycated haemoglobin (HbA1c) levels in blood reflect the average blood glucose levels during the preceding 2-3 months, corresponding to the average half-life of erythrocytes. Therefore, HbA1c levels provide a more reliable indication of glycemia over a longer period of time as compared to a single blood glucose measurement; in fact, HbA1c is used in clinical practice for diagnosis and prognosis of diabetes mellitus [27]. In line with the blood glucose data, anle138b treatment significantly reduced the mean blood HbA1c levels of *hIAPP Ob/Ob* mice, at 32 wk from 77 ± 3.3 mM/M (untreated) to 41 ± 5.3 mM/M (anle138b-treated) (*p* < 0.01); whereas in *Ob/Ob* mice HbA1c levels were not affected by anle138b treatment, being around 34 mM/M at 32 wk (Fig. 1e,f). Moreover, in treated *hIAPP Ob/Ob* mice blood glucose and blood HbA1c levels correlated significantly, while in non-treated *hIAPP Ob/Ob* mice and in the *Ob/Ob* mice (treated and non-treated) this correlation was not significant (Fig. 1g,h). Importantly, only in the *hIAPP Ob/Ob* mice treated with anle138b, was there a significant inverse correlation between fasting plasma insulin and blood glucose levels at 16 (*p* = 0.01), and 23 wk (*p* = 0.007); this correlation was borderline-significant at 32 wk (*p* = 0.07). On the other hand, no significant correlations were observed at all in the non-treated mice (*p* > 0. 57) (Fig. 1i), indicating that anle138b effectively restored the insulin-glucose relationship. Overall, these findings indicate that anle138b efficiently reduces blood glucose levels in diabetic *hIAPP Ob/Ob* mice and that this is mediated, at least partly, via beneficial effects on insulin production/secretion.

### 2.2 Anle138b affects insulin sensitivity in hIAPP Ob/Ob mice

Since beta cell function is related to insulin sensitivity, we next investigated whether anle138b influences insulin sensitivity by performing insulin sensitivity tests (ISTs). The relative blood glucose levels after intraperitoneal insulin injection, i.e. glucose levels relative to the level immediately prior to the insulin injection, were significantly lower in the anle138b-treated *hIAPP Ob/Ob* mice only at 8 wk (at the last timepoint, 150 minutes). At 24 wk there were no differences in relative glucose levels, whilst at 34 wk the relative blood glucose levels were actually higher in the anle138b-treated *hIAPP Ob/Ob* mice (Fig. 2a-c). In *Ob/Ob* mice, the relative glucose levels in ISTs were not changed by anle138b (Fig. 2d,e). In agreement with the relative glucose levels, the area under the curve (AUC) of the IST graph was reduced at 8 wk and increased at 34 wk in the anle138b-treated *hIAPP Ob/Ob* mice, while the AUC was not affected in the anle138b-treated *Ob/Ob* mice (Fig. 2g,h). However, these data are difficult to interpret as increased/decreased insulin sensitivity, because of the large differences in the basal/starting glucose levels of the treated vs the non-treated mice (Fig. S2a-c). Indeed, the basal/starting glucose levels of anle138b-treated mice were significantly lower at both 24 wk and 34 wk (49.6% and 61.% lower than in non-treated *hIAPP Ob/Ob* mice, respectively). Conversely, In *Ob/Ob* mice, insulin administration did not affect blood glucose levels differently in anle138b treated vs non-treated mice at any timepoint (Fig. 2d,e and Fig. S2d,e). Next, we evaluated glucose tolerance in *Ob/Ob* mice (glucose tolerance tests were not performed in the *hIAPP Ob/Ob* mice because of their severe hyperglycemia). Glucose injections to *Ob/Ob* mice increased blood glucose concentrations comparably in anle138b-treated vs non-treated mice (Fig. 2f and Fig. S2f), indicating that anle138b does not affect glucose tolerance in *Ob/Ob* mice. Overall, the data suggest changes of insulin sensitivity by anle138b in *hIAPP Ob/Ob* mice, but not in *Ob/Ob* mice.

**Figure 2.**
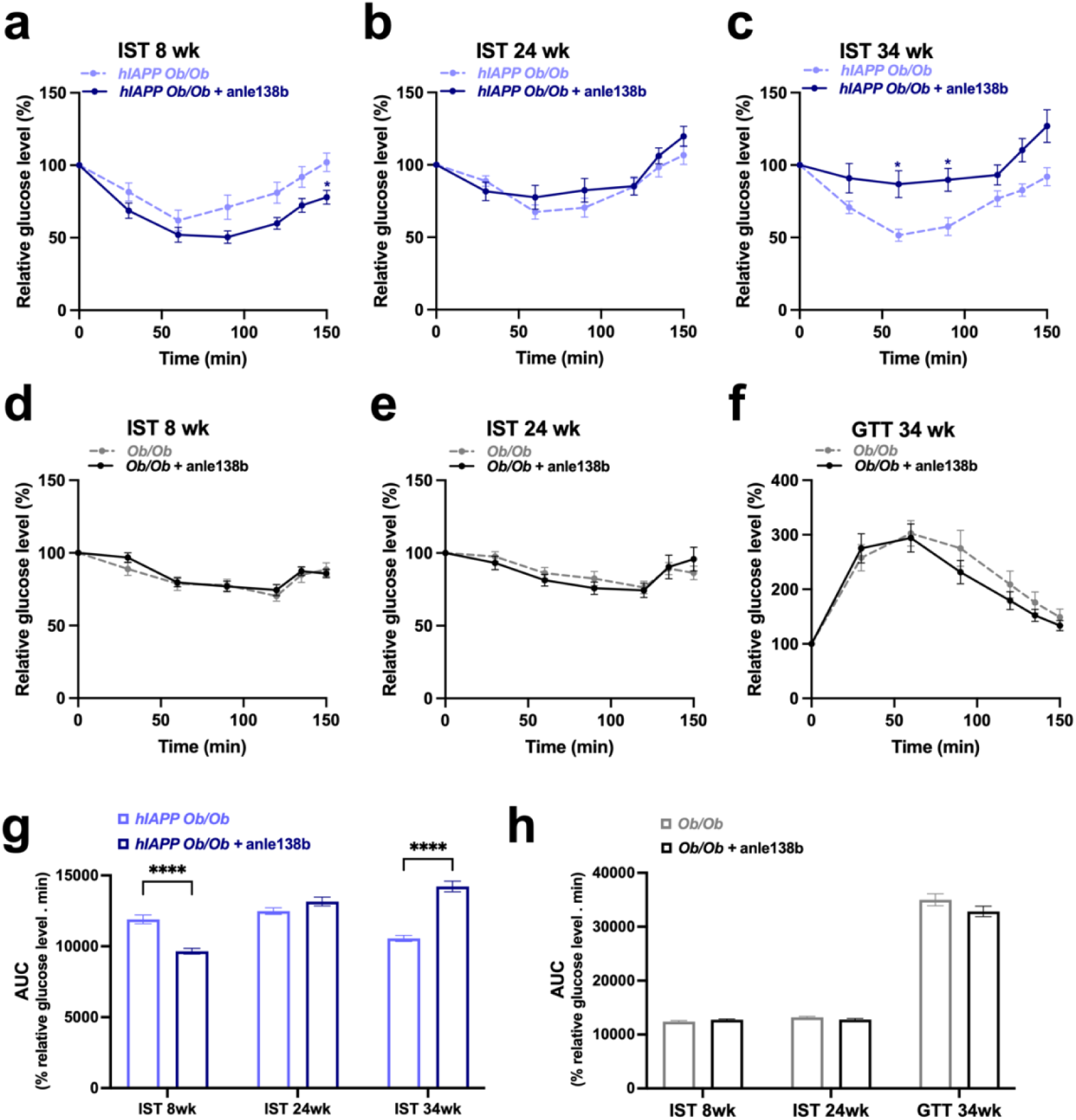
Insulin sensitivity test in *hIAPP Ob/Ob* and *Ob/Ob* mice, with and without anle138b treatment. (**a-c**) Insulin sensitivity test (IST) of *hIAPP Ob/Ob* mice after 8, 24 and 34 weeks of treatment, respectively. (**d,e**) IST of *Ob/Ob* mice after 8 and 24 weeks of treatment, respectively. (**f**) glucose tolerance test (GTT) of *Ob/Ob* mice after 34 weeks of treatment (*n* = 13-18 per group for the *Ob/Ob* mice and *n* = 12-15 per group for the *hIAPP Ob/Ob* mice at each time point). Insulin sensitivity and glucose tolerance were assessed by intraperitoneal injection of insulin (3 units/kg in *Ob/Ob* mice at 8 weeks, 4 units/kg in *Ob/Ob* mice at 24 weeks and 2.25 units/kg in *hIAPP Ob/Ob* mice at 8, 24 and 34 weeks) or glucose (0.8 gr/kg), respectively, after 6 h of fasting from 7 am onwards. Direct blood glucose measurements were performed immediately before injection and over a period of 150 min after injection. Blood glucose levels are indicated relative to the level at t=0 min. (**g,h**) Area under the curve (AUC, in % relative glucose level.min) for the graphs of panels **a-c** and **d-f**, respectively. All experiments were performed in male mice. (**a-h**) Data are presented as mean ± SEM; (**a-f**) Two-way ANOVA with Sidak’s test; \**p* < 0.05 for anle138b-treated vs untreated mice. Also see Figure S2. (**g,h**) unpaired two-way Welch’s t-test; \*\*\*\**p* < 0.0001 for anle138b-treated vs untreated mice.

### 2.3 Anle138b increases pancreatic islet number and beta cell mass in hIAPP Ob/Ob mice

Next, we assessed whether anle138b affects histopathological features of T2DM by examining pancreatic sections of the *hIAPP Ob/Ob* mice. The average wet-weight of the pancreas was indistinguishable between the treated and non-treated mice (154 ± 40 mg vs 166 ± 47 mg; *p* = 0.53). Amyloid was quantified in pancreatic sections stained with the amyloid-specific dye Congo red (Fig. 3a,b). Both the average percentage of islets containing amyloid (amyloid prevalence, Fig. 3c) and the average percentage of islet area consisting of amyloid (amyloid severity, Fig. 3d) were not significantly lower in *hIAPP Ob/Ob* mice treated with anle138b (*p* = 0.29 and *p* = 0.08, respectively). Remarkably, however, the number of islets was higher by 54% in pancreas of *hIAPP Ob/Ob* mice fed with anle138b (*p =* 0.005; Fig. 4a). In the *hIAPP Ob/Ob* mice treated with anle138b, islet number also correlated positively with fasting plasma insulin levels (Fig. 4b), and negatively with fasting blood glucose levels (Fig. 4c); importantly, these correlations were not observed in the non-treated *hIAPP Ob/Ob* mice. In notable agreement with the higher islet number, the total pancreatic beta cell mass (insulin-positive cell mass) was 58% bigger in the treated *hIAPP Ob/Ob* mice compared to the non-treated mice (*p* = 0.038; Fig. 4d-f).

**Figure 3.**
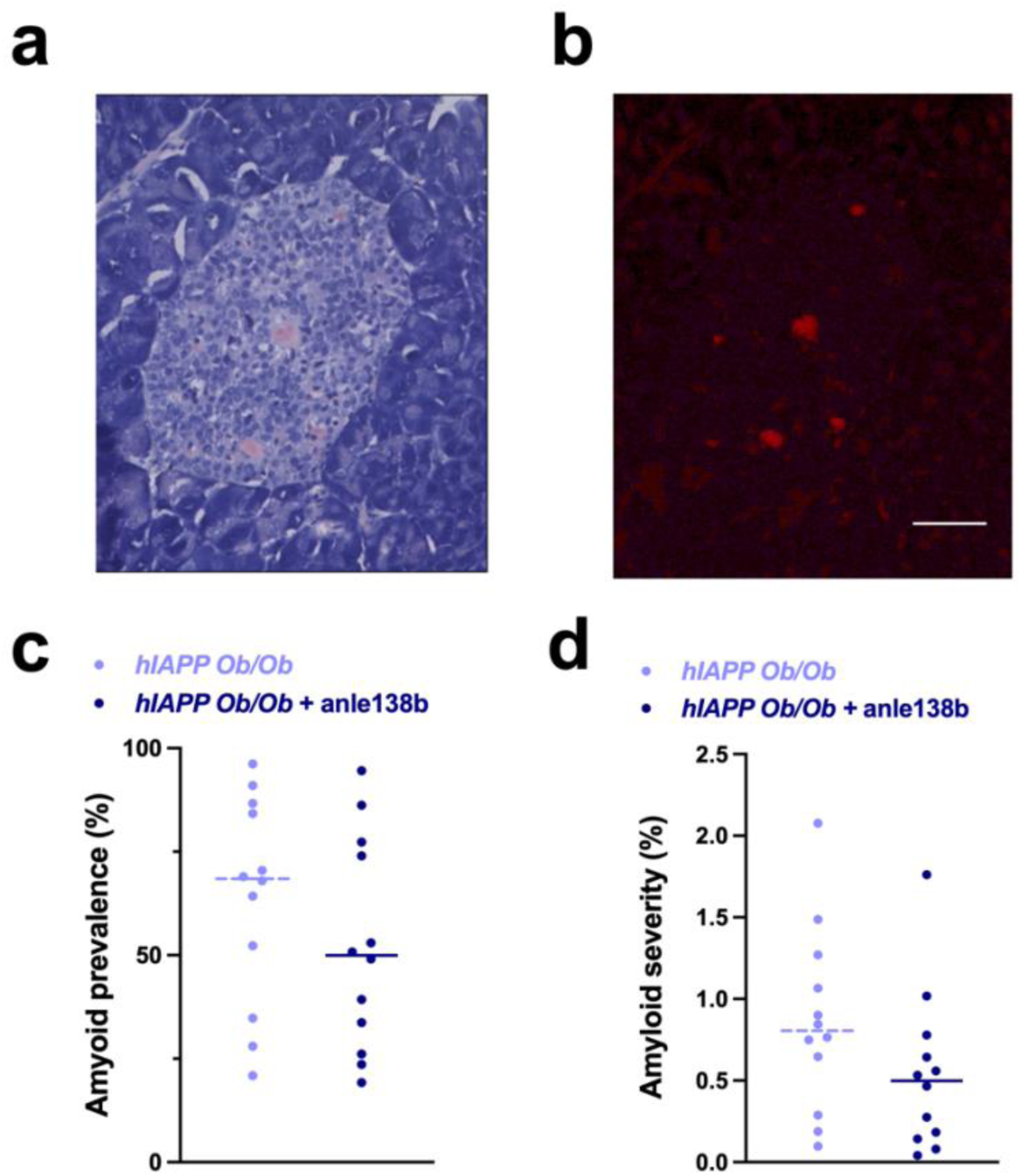
Anle138b does not reduce pancreatic islet amyloid load in *hIAPP Ob/Ob* mice. (**a,b**) Images of a section of a pancreatic islet from a *hIAPP Ob/Ob* mouse, stained with Congo red. Scale bar: 20 µm. (**a**) Brightfield image with pink color indicating amyloid deposits: islet amyloid; (**b)** Same islet section as in (**a)**, upon exposure to red light; the Congo red-positive amyloid deposits reveal a fluorescence which was quantified for each individual islet. (**c**) Islet amyloid prevalence (% of islets containing amyloid) and (**d**) islet amyloid severity (% of cross-sectional islet area being amyloid-positive). All experiments were performed in male mice (*n* = 12 per group). (**c,d**) The data are presented as the individual values and the median; Mann-Whitney test for anle138b-treated vs untreated mice.

**Figure 4.**
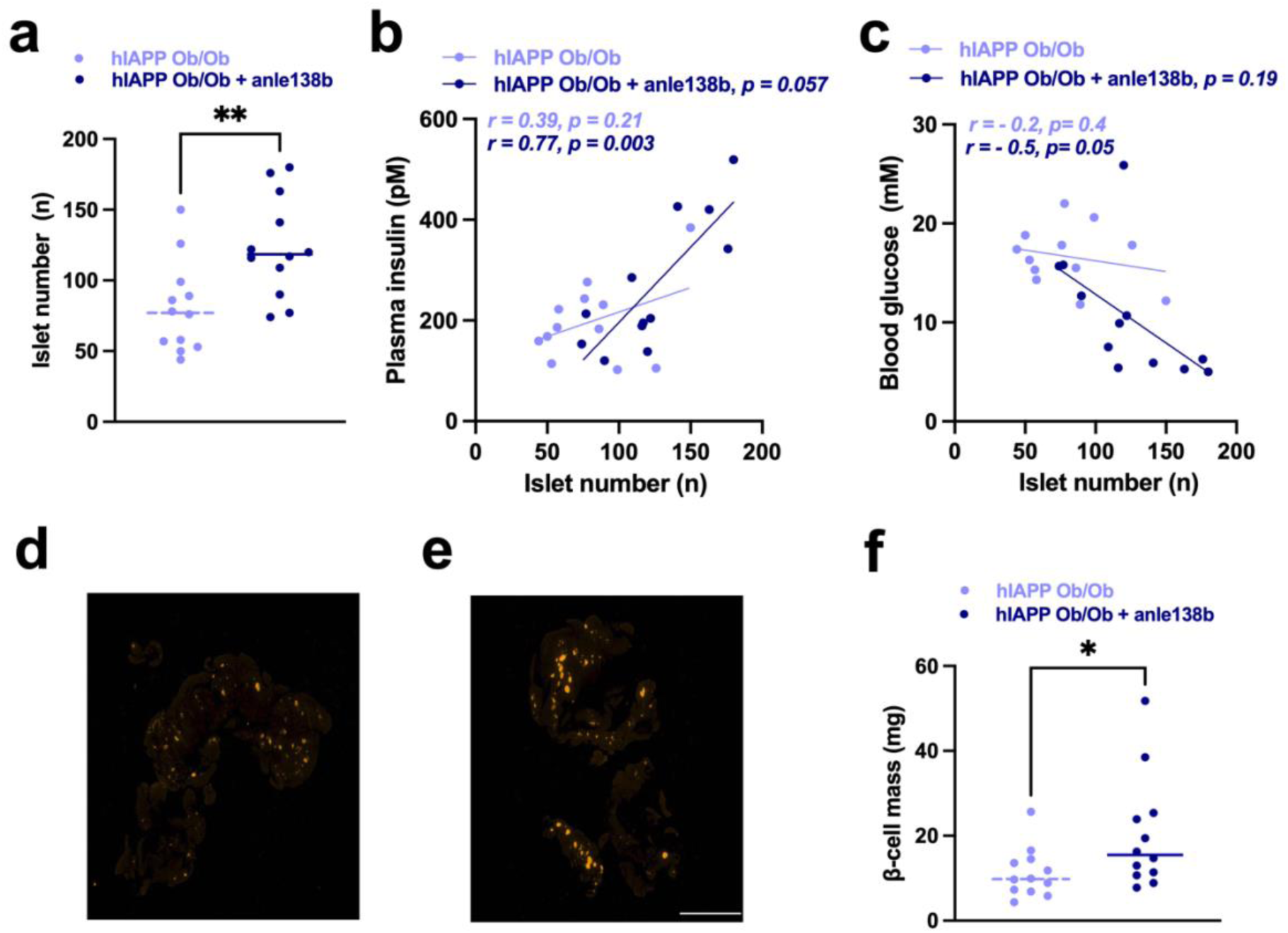
Anle138b increases pancreatic islet function and beta cell mass in *hIAPP Ob/Ob* mice. **(a)** Islet number and **(b,c)** correlations between islet number and metabolic parameters after 36 wk of treatment. All experiments were performed in male mice (*n* = 12 per group). **(a)** The data are presented as the individual values and the median; Mann-Whitney test for anle138b-treated vs untreated mice; \*\**p* < 0.01 vs *hIAPP Ob/Ob* mice without anle138b. **(b,c)** Pearson correlation test; correlation coefficient (r) and two-tailed *p*-value are indicated for both the treated and the untreated mice; *p*-value for linear regression analysis is indicated at the top of the figures. **(d,e)** Images of a pancreatic section stained for insulin (yellow color indicates insulin immunoreactivity) from a *hIAPP Ob/Ob* mouse, non-treated and treated with anle138b,respectively. Scale bar: 2 mm. **(f)** Quantification of total beta cell mass. The data are presented as the individual values and the median (*n* = 12 per group); Mann-Whitney test; \**p* < 0.05, vs *hIAPP Ob/Ob* mice without anle138b.

Overall, these data indicate that although anle138b reduces hyperglycemia in *hIAPP Ob/Ob* mice, the amyloid load of the pancreatic islets *per se* does not appear to be a strong/main determinant of islet function in these mice. Anle138b does, however, increase islet number and pancreatic beta cell mass, thus increasing the insulin-producing capacity of the pancreas.

### 2.4 Anle138b reduces hIAPP-induced damage to mitochondria in vitro

Mitochondrial impairment in pancreatic beta cells contributes to T2DM [28], and amyloid proteins, including hIAPP and amyloid-beta, cause damage to mitochondria [29]. Therefore, we investigated whether anle138b prevents hIAPP aggregate-induced damage to isolated mitochondria. Anle138b is membrane-permeable and thus has the ability to also reach intracellular targets such as mitochondria. Indeed, anle138b dramatically reduced hIAPP-induced cytochrome *c* release (CCR) from isolated mitochondria; when calculated as a percentage of maximal CCR induced by Triton X-100 detergent, anle138b decreased CCR from 53.2 ± 2.1% for mitochondria exposed to hIAPP aggregates to 19.3 ± 1.8% for mitochondria incubated with hIAPP + anle138b (*p* < 0.001); thus implying a 63.7% reduction of hIAPP-induced CCR as a result of anle138b treatment (Fig. 5a). Further, anle138b almost completely prevented mitochondrial swelling, from an average hydrodynamic diameter of 2,084 ± 236 nm for hIAPP-treated mitochondria to 1,146 ± 137 nm for mitochondria incubated with hIAPP + anle138b (*p* < 0.01); with control mitochondria having an average diameter of 1,065 ± 28 nm; this signifies a 92% decrease in mitochondrial swelling as a result of anle138b treatment (Fig. 5b). To gain further insights into how anle138b affects hIAPP-induced mitochondrial damage, we also evaluated changes in the mitochondrial membrane potential (ΔΨm), which reflects the integrity of the inner mitochondrial membrane. Preformed hIAPP protofibrillar aggregates reduced the ΔΨm of isolated mitochondria from a mean of 33,231 RFU to 21,916 RFU (*p* <0.001), as assessed by the fluorescent probe JC-1. Incubation with anle138b restored ΔΨm to a mean of 28,912 RFU (Fig. 5c), indicating that anle138b increased the ΔΨm by 62% when compared to the hIAPP-induced loss of mitochondrial membrane potential (*p* <0.001). Altogether, therefore, the *in vitro* data show that mitochondrial integrity affected by hIAPP is preserved by treatment with anle138b.

**Figure 5.**
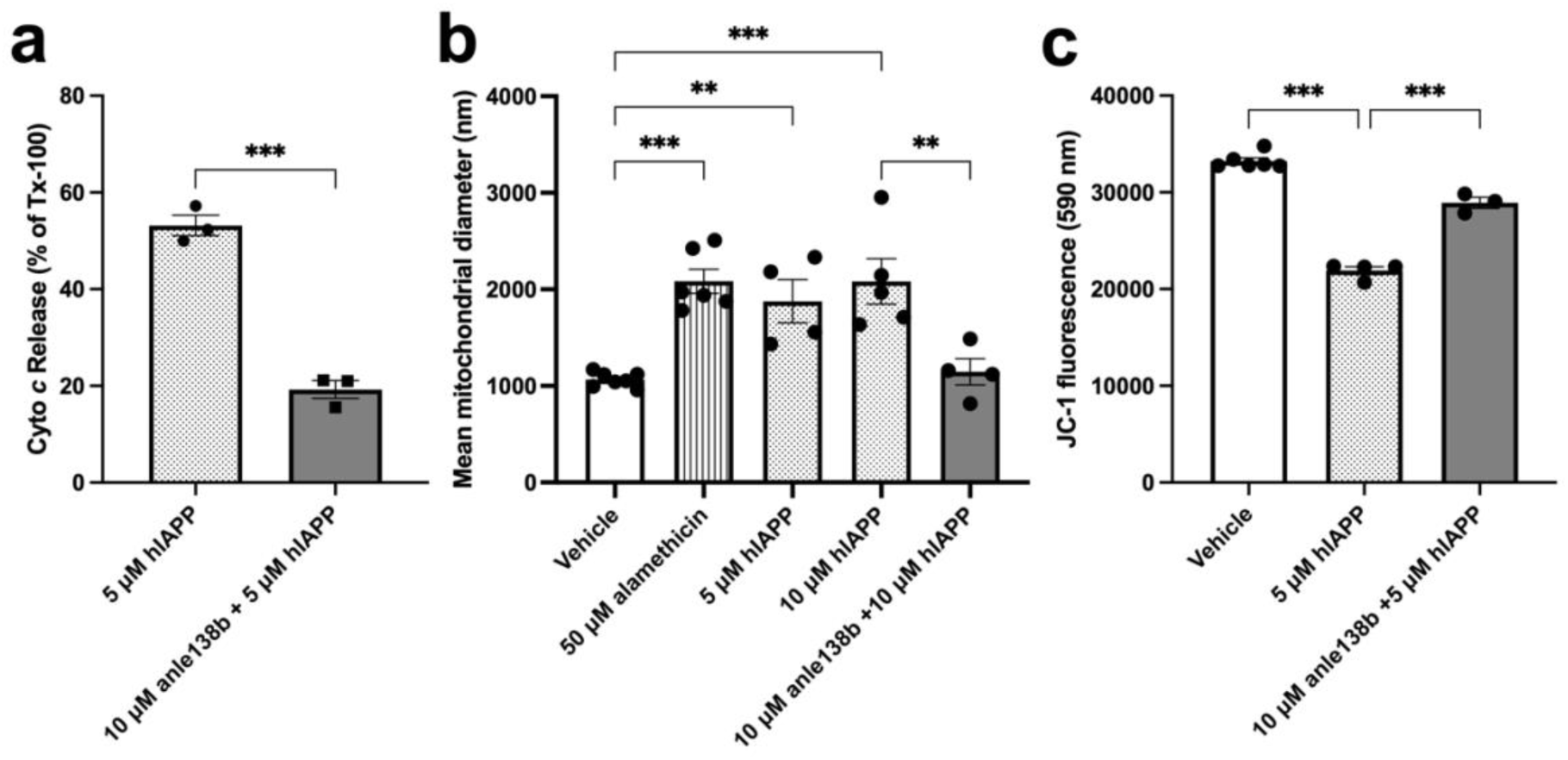
Anle138b has protective effects against hIAPP-induced mitochondrial damage. **(a**) Mitochondria from SH-SY5Y cells were incubated with hIAPP oligomers (5 μM) for 60 min, with or without pre-incubation with 10 μM anle138b for 10 min. The cytochrome *c* concentration (ng/ml) in the supernatant was quantified from hIAPP-treated mitochondria, with or without pretreatment with anle138b, and is indicated as the percentage of cytochrome *c* release by Triton X-100 (Tx-100). Bars indicate mean ± SEM (*n*=3; \*\*\**p* < 0.001; unpaired two-way Student’s t-test). (**b)** Fresh mitochondrial suspensions were exposed to swelling agent alamethicin, 5 μM or 10 μM hIAPP oligomers; or 10 μM hIAPP oligomers after pre-incubation with 10 μM anle138b for 10 min before addition of the oligomers. Bars indicate mean ± SEM of the average mitochondrial size measured by DLS. Comparisons were carried out by one-way ANOVA with Bonferroni’s posthoc correction, relative to vehicle treated mitochondria or to 10 μM hIAPP oligomers without anle138b (*n*=3-7; \**p* < 0.05, \*\*\**p* < 0.001). (**c**) End-point changes in the membrane potential (ΔΨm) of isolated mitochondria in mitochondrial buffer, monitored using the JC-1 dye, after 45 min in the absence and presence of 5 μM hIAPP oligomers, with or without pre-incubation with 10 μM anle138b for 10 min. Bars indicate mean ± SEM (*n*=3-6, \*\*\**p* < 0.001; one-way ANOVA with Bonferroni’s posthoc correction).

## 3. Discussion

In the present investigation, we assessed the potential therapeutic efficacy of the small-molecule amyloid oligomer modulator anle138b in T2DM, a common metabolic disease as well as a proteinopathy characterized by hIAPP aggregation and pancreatic islet amyloid formation [7,12]. The compound was administered through dietary admixture to transgenic *hIAPP Ob/Ob* mice, a model of severe T2DM, with treatment starting very early, at the time of weaning. Our study exclusively examined male mice, but it is expected that the findings are relevant also for female mice, since female hIAPP transgenic mice also develop islet amyloid and T2DM-related metabolic defects [13,14].

Oral administration of anle138b for 8 months resulted in a considerable improvement of glucose homeostasis, with a significant reduction of the basal blood glucose levels, both after overnight fasting and after a short fasting period (6 hr). As in clinical practice of diabetes mellitus [27], glycated hemoglobin (HbA1c) has been proven to be a reliable indicator of blood glucose regulation in diabetic mice [30]. In addition to the lowered blood glucose levels, we found that also the HbA1c levels were significantly reduced in the anle138b-treated *hIAPP Ob/Ob* mice. Moreover, anle138b restored the correlation between blood insulin and glucose levels in *hIAPP Ob/Ob* mice, implying that this compound reduces hyperglycemia by improving insulin production/secretion from the pancreatic islets. In the pancreas, anle138b treatment did not significantly reduce the islet amyloid load, suggesting that the beneficial metabolic actions of this compound are likely mediated through effects on smaller, prefibrillar aggregates of hIAPP. In the pancreas, anle138b treatment significantly increased the number of islets and the total beta cell mass. An increase in the pancreatic beta cell mass might also explain why at greater ages (Fig. 2c), the relative decrease in blood glucose levels during the ISTs became lower in anle138b-treated vs non-treated *hIAPP Ob/Ob* mice even though the same dose of insulin was injected. This indicates that the treated mice were less in need of additional (i.e. injected) insulin in conditions where the basal blood glucose levels had already been reduced substantially, suggesting that anle138b treatment has caused a reset of glucose homeostasis. This is also shown by the finding that hIAPP/ObOb mice treated with anle138b for 8 months had significantly reduced blood glucose levels whereas their insulin levels were similar to those of non-treated hIAPP/ObOb mice, indicating improved glucose homeostasis. Significantly, bodyweight and food intake were not affected by anle138b in *hIAPP Ob/Ob* mice, suggesting that this compound does not affect caloric intake and global metabolism.

Contrary to the *hIAPP Ob/Ob* mice, anle138b did not influence glycemia in non-transgenic *Ob/Ob* mice. Also insulin sensitivity and glucose tolerance were not affected by anle138b in these *Ob/Ob* mice which lack hIAPP. This implicates that the improved glucose homeostasis in anle138b-treated *hIAPP Ob/Ob* mice is due to effects of anle138b on hIAPP-mediated actions.

Interestingly, it was recently shown that anle138b treatment of male wildtype mice with insulin resistance (induced by a high-fat-diet [HFD]) caused improvements of glycemia and insulin sensitivity [31], indicating that anle138b can also have hIAPP-independent effects on glucose metabolism. In this HFD model, basal insulin levels are approximately 2-fold elevated as compared to untreated wildtype controls, whereas in the *Ob/Ob* and *hIAPP Ob/Ob* models basal insulin levels are around 10-fold and 5-fold higher, respectively, than in wildtype controls [14], thus indicating a more severe insulin resistance in our models. Nonetheless, both hIAPP-dependent and -independent effects of anle138b are expected to contribute to ameliorating the diabetic phenotype in humans with T2DM.

There are several other T2DM rodent models that also rely on hIAPP aggregation, with each model having its specific characteristics and limitations [15]. Notably, the *hIAPP Ob/Ob* model of T2DM used in this work recapitulates many features of the metabolic syndrome observed in human T2DM, including: obesity, dyslipidemia, insulin resistance, hyperinsulinemia, pancreatic islet amyloid deposits and beta cell failure [14]. The early development of hyperglycemia, at 4 wk of age, and the high blood glucose levels observed in these mice demonstrate the development of severe diabetes in this model [14]. The fact that anle138b can significantly reduce hyperglycemia even in this “aggressive” mouse model of severe T2DM robustly demonstrates the strong potential of this compound for T2DM.

The hIAPP-dependent, therapeutic effects of anle138b in a severe model of T2DM reinforce the notion, and previous studies indicating, that anle138b affects amyloid disorders via modulating pathogenic protein aggregation [21–23]. This is in contrast to other (postulated) mechanisms of anti-diabetic therapy; e.g. the autophagy inhibitor MSL-7 showed anti-diabetic effects in obese mice both with and without expression of hIAPP [32]. Indeed, our *in vitro* studies revealed that anle138b mitigated mitochondrial dysfunction caused by preformed hIAPP aggregates/oligomers. These findings are consistent with previous reports that demonstrated the ability of anle138b to reduce the impact of other amyloid proteins, i.e. α-synuclein, tau and amyloid-beta, on mitochondria [33]. Since mitochondria are intrinsically related to beta cell function/ insulin secretion [34], our collective experimental results therefore support the notion that hIAPP aggregation contributes to T2DM development through mitochondrial dysfunction [7,16]. Indeed, IAPP aggregation has previously been associated with mitochondrial dysfunction, in the form of oxidative stress, reduced mitochondrial glucose-stimulated respiration, an inhibition of ATP-coupled respiration, and, consequently, increased beta cell death, as also recently shown for isolated human islets as well as hIAPP transgenic mouse islets [35]. Hence, we put forward the notion that treatment with anle138b significantly suppresses hIAPP-induced beta cell pathology, as also indicated by the increased islet number and increased beta cell mass in anle138b-treated *hIAPP Ob/Ob* mice, and can restore glycemic control.

As stated previously, accumulation and aggregation of misfolded proteins are important factors in the development and progression of several neurodegenerative diseases [36]. Consequently, the use of agents/therapies capable of preventing and/or inhibiting pathological protein aggregation is a promising approach for slowing the development and progression of these diseases [37]. Also for human T2DM, which can be classified as a protein misfolding disease or proteinopathy, there is a great interest in strategies to prevent islet amyloid formation and its toxicity, and several therapeutic approaches in this field are currently under development [38,39]. For instance, IAPP protofibril-specific antibodies have shown potential in T2DM rodent models [18,19]. Small molecules, however, have numerous advantages compared to antibodies, including the potential for oral dosing and the ability to reach intracellular targets (such as mitochondria) by various mechanisms, mainly via passive diffusion in the case of lipophilic compounds like anle138b [20,38,39]. These are important considerations, particularly in the case of islet amyloidosis, since increasing evidence in rodent models shows that both extracellular and intracellular formation of toxic hIAPP aggregates can activate multiple overlapping pathological cellular signaling mechanisms leading to beta cell toxicity [11, 40, 41].

In conclusion, the beneficial effects of anle138b in the *hIAPP Ob/Ob* mouse model of severe T2DM attest to a strong therapeutic potential of this compound, and related DPP compounds, for T2DM. Anle138b preserved mitochondrial health against hIAPP aggregates *in vitro* while improving pancreatic islet function in the *hIAPP Ob/Ob* mouse model, allowing them to achieve lower blood glucose levels. Altogether, the data presented here strongly support the continued development of anle138b for a potential clinical application in T2DM and other proteinopathies, particularly since phase 1 studies of this compound demonstrated a well-tolerated safety profile in humans at potentially efficacious doses [25]. Finally, anle138b being able to modulate not only aggregation of hIAPP, but also that of α-synuclein, amyloid-beta and tau, could be beneficial not only for T2DM, but also ideally suited for the comorbidities observed between T2DM and PD or AD [42,43].

## 4. Materials and Methods

### 4.1 Mice

Transgenic mice overexpressing hIAPP in the pancreatic islet beta cells, under transcriptional control of a rat *Insulin 2* gene promoter, were generated as described previously [32] and back-crossed to C57Bl6J background. By cross-breeding the *hIAPP* transgenic line GG0018 with *Leptin-Ob* mice, developing obesity and mild hyperglycemia [44], the *hIAPP/Ob* line GG2653 was generated [14]. By selective breeding, a subline of GG2653 with the *Leptin-Ob* mutation but without the *hIAPP* transgene was obtained on the same genetic background (“non-transgenic *Ob*”). For this study, male *hIAPP/Ob* mice and non-transgenic *Ob mice* were treated and analyzed. Anle138b was administered to mice via dietary admixture with CRM(E) food (at a dose of 2 g anle138b/kg food), prepared by Ssniff Spezialdiäten GmbH (Soest, Germany). Mice from the control groups received CRM (E) food without anle138b (Ssniff Spezialdiäten GmbH, Soest, Germany). Further details on breeding, genotyping, housing and treatment of the mice are provided in the Supplementary Methods.

### 4.2 Bodyweight/food intake

Bodyweight was determined prior to each blood sampling, insulin sensitivity test (IST) and glucose tolerance test (GTT). Food intake was assessed after 6-7 months of treatment, by determining the decrease in weight of the food pellets placed on the grid of the cage on the first day of the measurement, and then again on the following 3 days (each day between 9-10 am). The daily decrease in weight on each of these 3 days was first averaged per cage, then averaged per mouse for the 2-3 mice sharing the same cage, and was expressed as g/mouse/day.

### 4.3 Blood sampling, IST, GTT

Immediately before start of the compound treatment (t=0, age 3-4 wk), blood was obtained by submandibular (cheek pouch) puncture (without anesthesia), after 4 h of fasting in the morning. After 16 and 32 wk of treatment, blood was obtained by submandibular puncture after overnight fasting. 30 µl of whole EDTA blood was taken for HbA1c determination and plasma was aliquoted for insulin- and IAPP measurements; samples were stored at -80⁰C until analyzed. Basal blood glucose measurements were performed during blood samplings at 0, 16, 23 and 32 wk, by directly assessing 1 drop of whole blood using an Accu Chek Aviva glucose meter (Roche Diagnostics, Germany).

In addition to the blood samplings described above, an intraperitoneal insulin sensitivity test (i.p. IST) was performed after 8 and 24 wk of treatment. In the *hIAPP Ob/Ob* mice, an additional IST was performed after 34 wk of treatment, whereas in the *Ob/Ob* mice an intraperitoneal glucose tolerance test (i.p. GTT) was performed after 34 wk of treatment GTT was not considered appropriate for the *hIAPP Ob/Ob* mice because of their high basal blood glucose levels.

All mice were killed by cervical dislocation after 34 wk (*Ob/Ob* mice, immediately after the GTT) or 36 wk (*hIAPP Ob/Ob* mice, after overnight fasting) of treatment. Several organs (including pancreas) were collected and stored for further analysis.

Further details on the IST and GTT, as well as methods used to determine blood glucose-, blood HbA1c-, plasma insulin- and plasma IAPP levels, are described in the Supplementary Methods.

### 4.4 Histological analyses

After pancreas collection, the wet weight was determined and the tissue was fixed in 10 ml of 10% buffered formalin (overnight at room temperature). The formalin-fixed tissue was dehydrated and embedded in paraffin blocks. For quantification of islet amyloid content and the pancreatic beta cell mass, sections of formalin-fixed paraffin embedded pancreas were cut at 3 different regions (“depths”) and stained with Congo red (Agilent Technologies, AR16192-2 Artisan Congo red stain kit)/haematoxylin, as described previously (14) or with an insulin antibody (Invitrogen, clone 4C3Y9, 1:200), respectively. The amyloid was detected by Congo red fluorescence upon illumination with red light; insulin-stained beta cells were detected using a fluorescently-labeled secondary antibody. Quantification of islet amyloid and beta cells was peformed as described in the Supplementary Methods.

### 4.5 Preparation of hIAPP oligomers for mitochondria experiments

1.4 mM stock solutions of hIAPP_1-37_ (Abcam, UK) were prepared by dissolving 1 mg of lyophilised peptide in 99.9% DMSO/0.1% TFA. Aliquots of hIAPP stock were kept in LoBind® epitubes (Eppendorf) and snap-frozen immediately in liquid nitrogen to prevent aggregation. All samples were stored at -80 °C. Aggregation of 5 μM fresh monomeric hIAPP into oligomers was carried out in a 96-well microtitre plate under shaking conditions, and oligomer characterization was performed as further described in the Supplementary Methods.

### 4.6 Mitochondrial assays

Mitochondria were isolated from SH-SY5Y cells (cloned subline of neuroblastoma from a female patient with a metastatic bone tumor) [45]. The SH-SY5Y cell line was newly obtained from the European Collection of Authenticated Cell Cultures (ECACC #94030304), and always tested negative in routine tests for mycoplasma contamination. Mitochondrial assays (swelling, cytochrome c release, and membrane potential) were all performed on freshly harvested mitochondria (0.125 mg/ml). The mitochondrial assays were performed as described previously [45] and in the Supplementary Methods.

### 4.7 Statistical analysis

Data were analysed using software R, tidyverse (DOI: 10.21105/joss.01686), and ggpubr and a *p*-value of ≤ 0.05 was considered to indicate a statistically significant difference/correlation. Normal distribution of data was tested using the Shapiro–Wilk test and QQ-plots. In case data were not normally distributed, non-parametric tests were used. Unpaired two-way t-test was used for pairwise comparisons of normally distributed data. One-way ANOVA with Bonferroni’s post hoc test and two-way ANOVA with Tukey’s or Sidak’s post hoc test were used for multiple comparisons. Mann–Whitney test (unpaired) was used for comparison of not-normally distributed data. Pearson correlation test was used for correlation analysis of two parameters with normally distributed data; linear regression was performed to examine if the slopes of the two regression lines are significantly different.

Graphs were prepared using GraphPad Prism 10 (CA, USA). Data are shown as means ± SEM and/or as individual datapoints with their median value. The statistical tests applied, *n-*values, correlation coefficient (*r*) and *p*-values are indicated in the figure legends.

### Study approval

The procedures concerning animal care, treatment and experimentation for this study were performed in accordance with the ARRIVE 2.0 guidelines for animal experimentation and with previous approval from the local Committee for Animal Experimentation of Utrecht University and University Medical Center Utrecht (DEC), the local Experimental Animal Welfare body (IVD, Workprotocol 323-6-01) and the national Central Authority for Scientific Procedures on Animals (CCD) (license number AVD115002015323).

## Supporting information

Supplemental methods plus supplemental figures

## Acknowledgments

We thank Anja van der Sar and Trudy Oosterveld-Romijn (GDL, Utrecht University, Utrecht, The Netherlands) for assistance with the mouse experiments; Inge Maitimu (Clinical Chemistry, University Medical Center Utrecht, The Netherlands) for the blood HbA1c, plasma insulin and plasma IAPP measurements; Domenico Castigliego (Pathology, University Medical Center Utrecht, The Netherlands) for scanning of the Congo red-stained pancreatic sections; Jens Brüning (Max Planck Institute for Metabolism Research, Cologne, Germany) for ongoing discussions during the research, and Andisheh Abedini (New York University, New York City, USA) for critical reading of the manuscript.

## Data availability

Values for all data points in graphs (Figures 1–5 and S1-S2) are reported in the Source Data XLS files. This study includes no data deposited in external depositories.

## Funding

This work was financially supported by MODAG GmbH (Project grant), the Max Planck Society (Institutional funding by a project grant), the Malta Council for Science & Technology (MCST) through the Research Excellence Programme (REP-2021-016), the Tertiary Education Scholarship Scheme (TESS) of the Ministry for Education and Employment of Malta (grant number MDSBP10-18) and the King Abdulaziz City for Science and Technology (KACST), Saudi Arabia (grant number 8091/10).

## Authors’ relationships and activities

A. G. and C. G. are co-founders of MODAG GmbH. A.G. is a full-time employee of MODAG. A. L. and S.R. are partly employed by MODAG and are beneficiaries of the phantom share program of MODAG. A.L., S.R., C.G. and A.G. are co-inventors of WO/2010/000372. Anle138b is licensed by Teva Pharmaceutical Industries Ltd (Tev-56286; Emrusolmin) and is in clinical development in collaboration with MODAG.

## Contribution statement

**Jo Höppener:** conceptualization, methodology, investigation, formal analysis, data curation, writing-original draft, writing- review & editing, visualization, supervision, project administration, funding acquisition. **Christian Griesinger:** conceptualization, writing-original draft, writing- review & editing, supervision, project administration, funding acquisition. **Niels Eijkelkamp:** writing- review & editing, supervision. **Armin Giese:** conceptualization, funding acquisition. **Sergey Ryazanov:** conceptualization, methodology, resources. **Neville Vassallo:** methodology, formal analysis, data curation, visualization, writing- review and editing, supervision. **Mohammed Albariqi:** investigation, formal analysis, data curation, writing-original draft, writing- review & editing, visualization, **Sanne Baauw**: investigation, formal analysis, data curation. **Sjors Fens:** investigation, formal analysis. **Sabine Versteeg:** investigation. **Raina Marie Seychell:** investigation. **Adam El Saghir:** investigation. **Lucie Khemtemourian:** investigation. **Nikolas Stathonikos:** methodology, software. **Andrei Leonov:** methodology, resources. **Hanneke Willemen:** investigation, data curation. **Bram Gerritsen:** formal analysis.

## Notes

### Summary of Updates

This version of the manuscript has been streamlined and updated for journal submission. Some text and data have been removed to focus on the manuscript's core findings; in addition some new data have been added

