## Supplemental methods plus supplemental figures for "Anle138b manifests potent anti-hyperglycemic activity in type 2 diabetic mice expressing hIAPP"

***Supplementary Methods***

**Mice**

hIAPP transgenic founder mice on a C57BL6J/DBA2 background were back-crossed to C57Bl6J. Homozygous *hIAPP* mice (*hIAPP*$^{+/+}$) were obtained by intrabreeding hemizygous *hIAPP* mice (*hIAPP*$^{+/-}$) and selecting for homozygous offspring by quantitative Southern dot blot hybridization [26].

*Leptin-Ob* mice on a C57Bl/6J background were originally obtained from Harlan. Homozygous inactivation of the *Leptin* gene in these mice causes obesity and mild hyperglycemia [44]. By cross-breeding the *hIAPP* transgenic line GG0018 with *Leptin-Ob* mice, the *hIAPP/Ob* line GG2653 was generated [14]. By selective breeding, a subline of GG2653 with the *Leptin-Ob* mutation but without the *hIAPP* transgene was obtained on the same genetic background ("non-transgenic Ob"). Presence of the inactivating missense mutation in the *Leptin* gene from *Leptin-Ob* mice was detected by PCR of ear DNA with primers flanking the point mutation (Ob1: 5' TGC CTT CCC AAA ATG TGC TGC 3' and Ob2: 5' CAT TCA GGG CTA ACA TCC ACC 3') and subsequent DdeI digestion of the 250 bp PCR product, which reveals the *Leptin-Ob* point mutation. Presence of the *hIAPP* transgene was proven by PCR with a forward primer in the rat *Insulin 2* gene promoter (RIPRev3F: 5' GAGATGGAGACAGCTGGCTC 3') and a reverse primer in exon 1 of the h*IAPP* gene (8910$^{+}$: 5' GTCAGCAATATCAGCAAATGCTTCTG 3'), generating a product of approximately 700 bp in *hIAPP* transgenic mice only. Homozygosity for the *hIAPP* transgene in the *hIAPP/Ob* line was assessed using quantitative Southern dot blot hybridization [26]. In mice used for breeding *hIAPP Ob/Ob* offspring for this study, hIAPP homozygosity was confirmed by previous test matings with non-transgenic mice, yielding only hIAPP transgenic offspring.

All the mice were housed and bred in polypropylene cages containing hardwood bedding and environmental enrichment (cardboard boxes and tissues) and the mice were maintained in airconditioned rooms at 20-22 °C on a 12 h light/dark cycle in the Animal house of Utrecht University and the UMC Utrecht (GDL, which has an AADDCC license accreditation). Food and water were provided ad libitum.

For this study, only male mice from the non-transgenic Ob and the homozygous hIAPP transgenic Ob sublines were used. Mice were genotyped at an age of 2-3 wk: pups were individually marked by ear cuts and the ear tissue removed was used for DNA analysis (PCR and DdeI digestion). Non-transgenic obese (*Ob/Ob*) as well as homozygous *hIAPP* transgenic obese (*hIAPP Ob/Ob*) males were treated with anle138b for 34-36 wk from weaning (age 3-4 wk) onwards. Male *Ob/Ob* and *hIAPP Ob/Ob* littermates were also used in the control groups, which received CRM(E) food without anle138b. *Ob/Ob* and *hIAPP Ob/Ob* mice were randomly assigned to each of the two treatment groups (with or without anle138b) by coin toss; when possible, pups from the same litters were split into the two treatment groups. Since anle138b had not been used previously in diabetic mice, and effect size was thus unknown, sample size calculation could not be performed *a priori.* For each treatment group 15 (*hIAPP Ob/Ob*) or 18 (*Ob/Ob*) mice entered the study (total 66 mice); with 2-4 mice sharing a cage. In case of suspicion of unexpected discomfort, based on abnormal behavior, posture, mobility or other clinical sings, bodyweight was determined daily. If bodyweight of a mouse decreased > 15% within 1 week, this animal was euthanized. Because of their obesity, *(hIAPP)Ob/Ob* mice from age 7 months onwards sometimes cannot get on their feet again after falling over. Therefore these mice were checked for this daily (also in the weekends) and if this happened for a mouse >3 times in a week , the humane endpoint was reached and this mouse was euthanized. For the hIAPP Ob/Ob mice, 3 animals died prematurely in both the anle138b-treated and non-treated groups. For the *Ob/Ob* mice, 2 animals died prematurely in the non-treated group only. Both the animal care takers of the GDL and the investigators performing the experiments and analyses were blinded to the treatment; the different foods were coded before arrival at the GDL and these codes were not cracked until all data had been collected and analyzed.

**Blood sampling, IST, GTT**

IST: mice were fasted for 6 h from 7 am onwards and injected intraperitoneally with insulin at a dose of 3 U/kg (*Ob/Ob* mice at 8 wk), 4 U/kg (*Ob/Ob* mice at 24 wk) or 2.25 U/kg (*hIAPP Ob/Ob* mice at 8, 24 and 34 wk). Human insulin intrarapid (Sanofi, France) was diluted with 0.9% NaCl to a concentration of 0.4 mU/µl, which was used for the i.p. injections. Multiple bloodsamples until 150 minutes after insulin injection were obtained by tail-tip blood sampling in unrestrained mice and 1 drop of blood was used for direct blood glucose measurements using an Accu Chek Aviva glucose meter (Roche Diagnostics). When blood glucose levels exceeded the upper limit of the glucose meter (33,3 mM) in *hIAPP Ob/Ob* mice, a second bloodsample was taken immediately, diluted with 0.9% NaCl and measured. The read-out of these diluted blood samples was corrected accordingly.

GTT: mice were fasted for 6 h from 7 am onwards and injected intraperitoneally with glucose at a dose of 0.8 g/kg. A 20% glucose solution in water was diluted 1:1 with 0.9% NaCl to a concentration of 10%; from this solution, 8 µl was injected per g bodyweight, corresponding to a dose of 0.8 g glucose per kg bodyweight. Multiple bloodsamples until 150 minutes after glucose injection were obtained by tail-tip blood sampling in unrestrained mice and 1 drop of blood was used for direct blood glucose measurements using an Accu Chek Aviva glucose meter (Roche Diagnostics).

For the *Ob/Ob* mice, the data from 1 mouse for the first IST (at 8 wk) and from 5 mice for the GTT (at 34 wk, 3 non-treated mice and 2 treated mice) were not included because the read-out of the glucose-meter exceeded the upper limit).

Blood HbA1c (mM glycated Hemoglobin/M total Hemoglobin) was determined in 30 µl of full (EDTA) blood by a direct whole blood enzymatic assay (Abbott Diagnostics, USA, product number 3L82-21)

Plasma insulin was determined using a rat insulin RIA from Merck-Millipore (cat # RI-13K) cross-reacting with mouse insulin.

Plasma IAPP was determined using a human IAPP (Amylin) ELISA from Merck-Millipore (cat # EZHA-52K) cross-reacting with mouse IAPP.

**Histological analyses**

For islet amyloid quantification, Congo red stained pancreatic sections of 5 µm were scanned using a Hamamatsu nanozoomer 2.0 RS, both in brightfield as well as in fluorescence mode using a TRITC filter (red channel); the brightfield- and fluorescence scans of each slide had an identical size. The resulting scans were annotated manually to select the islets (using the bright-field scans) and binary masks were generated to calculate the total cross-sectional area of islets. Using the red channel of the fluorescence scans, the islets were localized and by thresholding the fluorescence signal (using a predefined threshold), the area of amyloid in each individual islet was calculated over the total islet area. The total number of pancreatic islets, the average percentage of amyloid-positive islets ("amyloid prevalence") and the average percentage of amyloid-positive islet area ("amyloid severity") were determined for each mouse, by using the cumulative data from the 3 sections analyzed for each mouse.

For quantification of the beta cell mass, pancreatic sections of 3 µm were stained with a rabbit anti-human insulin antibody (Invitrogen, clone 4C3Y9, 1:200) and a goat anti-rabbit Alexa Fluor 555 secondary antibody (Invitrogen, 1:200). The sections were mounted with Vectashield hardset with DAPI (Vector). The slides were scanned using a Hamamatsu nanozoomer 2.0 RS, both in bright-field as well as in fluorescence mode using a red filter. The total cross-sectional tissue area on each slide was quantified in QuPath (version 0.4.4) using the DAPI staining. The cross-sectional insulin-positive pancreas area on each slide was determined in Qupath using the immunofluorescence staining. The percentage of insulin-positive pancreas area (averaged from the 3 sections at the different regions) was multiplied by the wet weight of the total pancreas (determined at the time of organ collection) to quantify the total pancreatic beta cell mass of each mouse in mg.

**Preparation of hIAPP oligomers for mitochondria experiments**

hIAPP aggregation was carried out in a 96-well microtitre plate, by incubating 5 μM fresh monomeric hIAPP in 10 mM MOPS (4-morpholinepropanesulfonic acid)/Tris, pH 7.4, buffer in a total volume of 200 μl. The plate was kept under shaking conditions (450 rpm) in a Comfort Thermomixer® (Eppendorf) at 37 °C for 45 min until oligomer formation. The latter was confirmed by (i) particle sizing using DLS (< 60 nm); (ii) positive reactivity to the anti-oligomer antibody A11 [46] (Millipore, AB9234; 1:2000) in an immunoblot dot test as described previously [47]); and (iii) using the fluorescence probe DCVJ [9-(2,2-dicyanovinyl) julodine] (Sigma-Aldrich, Cat#72335), which exhibits high sensitivity for early protein misfolding aggregates in the lag phase [48], and Thioflavin-T (ThT) (Sigma-Aldrich, T3516), which detects the cross-β structure of amyloid fibrils [49]. For the DCVJ and ThT assays, 5 μM DCVJ and 10 μM ThT, respectively, were added to the hIAPP samples and readings taken in a BioTeK FLx800 fluorescence reader with excitation at wavelength/filter 440/30 nm and emission captured at wavelength/filter 500/27 nm (DCVJ) or 485/20 nm (ThT). Fresh, non-aggregated hIAPP directly from stock (DCVJ-negative and ThT-negative) and 2-hr aggregated fibrillar hIAPP (DCVJ-negative and ThT-positive) were included for direct comparison with the oligomeric samples (DCVJ-positive and ThT-negative). The latter were either used immediately, or kept on ice and used within 1 h of preparation.

**Mitochondrial swelling assay**

Swelling assays on isolated mitochondria (0.125mg/ml) were performed at 25 °C in a low-volume disposable sizing cell, with readings taken every 10 min for 1 h using the multiple narrow modes analysis model of the ZS Xplorer® software [45]. Mitochondria were incubated without treatment, with hIAPP oligomers and with hIAPP oligomers + anle138b. In the latter condition, anle138b was incubated with mitochondria for 10 min prior to addition of the hIAPP oligomers. The pore-forming peptide alamethicin was used as a positive control agent for mitochondrial swelling.

**Mitochondrial cytochrome c release (CCR) assay**

A cytochrome c release (CCR) assay (R&D systems Quantikine® ELISA) was employed to measure direct mitochondrial membrane damage by hIAPP oligomers, as described previously [45]. Here, isolated mitochondria (40 µg) were incubated at 30°C for 60 min with hIAPP oligomers and with hIAPP oligomers + anle138b. In the latter condition, anle138b was incubated with mitochondria for 10 min prior to addition of the hIAPP oligomers. The final volume in this assay was 100 µl in 1× mitochondrial storage buffer (MITOISO2®, Sigma-Aldrich, Germany). The amount of cytochrome c present in the supernatant was quantified and results expressed as percentage of CCR induced by 1% (v/v) Triton X-100 detergent (Sigma-Aldrich, T8787) (theoretical maximium: 100%).

**Mitochondrial membrane potential (JC-1) assay**

The JC-1 assay (Sigma-Aldrich, CS0760) was performed as described previously [45] using 5 µg of isolated mitochondria at 30 °C, with fluorescence top readings taken every 5 min at wavelength/filter 485/20 nm for excitation and 590/10 nm for emission in a microplate reader (Tecan Infinite® 200 Pro). Mitochondria were incubated without treatment, with hIAPP oligomers and with hIAPP oligomers + anle138b. In the latter condition, anle138b was incubated with mitochondria for 10 min prior to addition of the hIAPP oligomers. In each experiment, the protonophore FCCP (5 μM) (Sigma-Aldrich, C2920) was used to induce complete collapse of the ΔΨm; this baseline fluorescence value in the presence of FCCP was subtracted from all mitochondrial sample values.

### Supplementary Figures

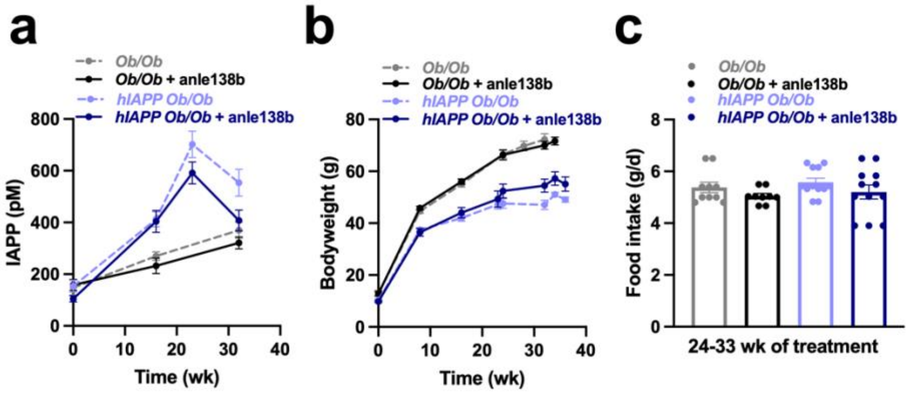


**Figure S1. Plasma IAPP levels, bodyweight and food intake of *Ob/Ob* and *hIAPP Ob/Ob* mice, with and without anle138b treatment.**

**(a)** Basal plasma IAPP levels of *Ob/Ob* and *hIAPP Ob/Ob* mice determined after 4 h fasting in the morning (t=0 weeks), after 6 h fasting in the morning (at 23 weeks) or after overnight fasting (at 16 and 32 weeks) (*n*= 17-18 per group for the *Ob/Ob* mice and *n* = 12-15 per group for the *hIAPP Ob/Ob* mice at each timepoint). **(b)** Bodyweight of *Ob/Ob* and *hIAPP Ob/Ob* mice determined after 4 h fasting in the morning (t=0 weeks), after 6 h fasting in the morning (at 8, 23, 24 and 34 weeks) or after overnight fasting (at 16, 32 and 36 weeks) (*n*= 16-18 per group for the *Ob/Ob* mice and *n* = 12-15 per group for the *hIAPP Ob/Ob* mice at each time point). **(c)** Daily food intake determined at 24-27 weeks of treatment for *Ob/Ob* mice (*n*= 9-10) and at 28-33 weeks of treatment for *hIAPP Ob/Ob* mice (*n*=12). Two-way ANOVA with Tukey's test; for anle138b-treated vs untreated mice.

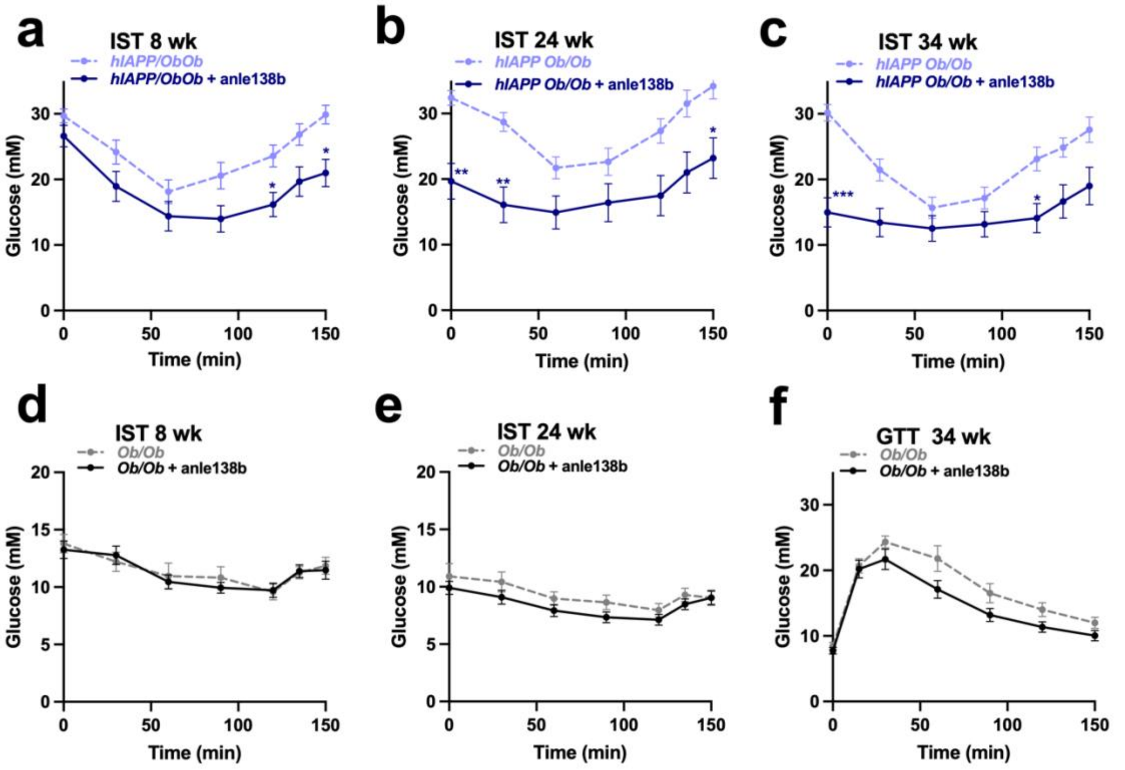


**Figure S2. Insulin sensitivity test of *hIAPP Ob/Ob* and *Ob/Ob* mice, with and without anle138b treatment.**

(**a-c**) Insulin sensitivity test (IST) of *hIAPP Ob/Ob* mice after 8, 24 and 34 wk of treatment, respectively. (**d,e**) IST of *Ob/Ob* mice after 8 and 24 wk of treatment, respectively ($n$ = 13-18 per group for the *Ob/Ob* mice and $n$ = 12-15 per group for the *hIAPP Ob/Ob* mice at each time point). (**f**) Glucose tolerance test (GTT) of *Ob/Ob* mice, after 34 wk of treatment ($n$ = 13-16 per group at each time point). Insulin sensitivity and glucose tolerance were assessed by intraperitoneal injection of insulin (3 units/kg in *Ob/Ob* mice at 8 wk, 4 units/kg in *Ob/Ob* mice at 24 wk and 2.25 units/kg in *hIAPP Ob/Ob* mice at 8, 24 and 34 wk) or glucose (0.8 gr/kg), respectively, after 6 h of fasting from 7 am onwards. Direct blood glucose measurements were performed immediately before injection and over a period of 150 min after injection. Data are presented as mean ± SEM; Two-way ANOVA with Sidak's test; *$p < 0.05$, **$p < 0.01$, ***$p < 0.001$ for anle138b-treated vs untreated mice.
